# Successful optimization of reconstruction parameters in structured illumination microscopy – a practical guide

**DOI:** 10.1101/402115

**Authors:** C. Karras, M. Smedh, R. Förster, H. Deschout, J. Fernandez-Rodriguez, R. Heintzmann

**Affiliations:** Leibniz Institute of Photonic Technology (IPHT), Jena, Germany; Centre of Cellular Imaging, Core Facilities, the Sahlgrenska Academy, Univ. of Gothenburg, Sweden; Institute of Physical Chemistry, Friedrich-Schiller-University (FSU) Jena, Germany

**Author notes:** Corresponding Author: Christian Karras.

## Abstract

The impact of the different reconstruction parameters in super-resolution structured illumination microscopy (SIM) onto artifacts is carefully analyzed. They comprise the Wiener filter parameter, an apodization function, zero-frequency suppression and modifications of the optical transfer function. A detailed investigation of the reconstructed image spectrum is concluded to be suitable for identifying artifacts. For this purpose, two samples, an artificial test slide and a more realistic biological system, were used to characterize the artifact classes and their correlation with the image spectra as well as the reconstruction parameters. In addition, a guideline for efficient parameter optimization is suggested and the implementation of the parameters in selected up-to-date processing packages (proprietary and open-source) is depicted.

## Introduction

To image fine features, large angels of light emitted by the sample have to be brought to interfere at the detector plane. However the range of acceptance angles of the objective sets a limit, leading to Abbe’s equation of resolution (1). The minimum feature size is represented as a grating possessing a period of

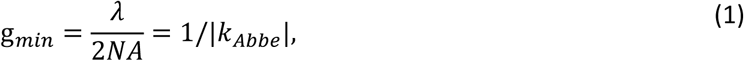

with *λ* being the wavelength of light in vacuum, *NA*=*n* sin(*α*) denoting the numerical aperture at refractive index of the embedding medium *n* and the acceptance angle of the objective *α*. |*k*_*Abbe*_| refers to the maximally transmitted spatial frequency. Fourier analysis allows to decompose a given object (or image) into a sum of waves. Each such composite wave has a specific period as measure for the feature size, i.e. a spatial frequency, intensity, phase and direction. The complete decomposition into such waves is known as image spectrum.

In fluorescence microscopy, one possibility of resolving finer structures than in (1) is to illuminate the sample with a structured (typically sinusoidal shaped stripes) excitation light pattern (2-4). Thereby, spatial object frequencies beyond 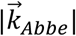 are down-modulated into the passband of the objective prior to detection. This technique, known as “structured illumination microscopy” (SIM), is superior in terms of photo toxicity and acquisition speed of super resolved images compared to other methods (5-10). Additionally SIM is not restricted to a particular cell staining. Thus it is considered one of the most suited super-resolution technologies for exploring dynamical processes in living cells (11-16).

The downside of SIM is its challenging image reconstruction, requiring a diligent experimental implementation and parameter optimization in order to avoid artifacts in the final image (17). Recent comprehensive protocols for successfully performing SIM experiments (18, 19), only marginally deal with the image reconstruction. Most published work on analyzing parameter-related artifacts in SIM, focusses on optimizing the estimate of the illumination patterns (20, 21). Properly choosing reconstruction parameters such as for Wiener filtering, apodization, zero-order suppression or adaption of the transfer function, still remains a challenging task for the end user of SIM. The present work aims to unravel the correlation between those parameters and typical artifacts in the final image. Additionally, a guideline for parameter optimization is suggested and the implementation of parameters in different available reconstruction software packages is compared.

## Experimental

Two different samples were used in order to study the impact of reconstruction parameters on the post-processing of structured illumination images: i) A photostable commercial sample (Argo-SIM slide, Argolight, France) consisting out of fluorescing double line pairs (spacing from 150 nm to 240 nm, λ_ex_ = 360 −550 nm, line thickness below 100 nm). ii) HeLa cells, fixed and processed for indirect immunofluorescence stain to reveal the endogenous Golgi-resident glycosylation enzyme, N-acetylgalactosaminyltransferase-T1 (GalNAc-T1) at the *trans* Golgi cisternae.

Structured illumination raw images were recorded using a commercial ELYRA PS.1(Carl Zeiss, Germany) microscope. Three and five grating orientations for the Argolight slide and for the GalNAc-T1 sample were recorded, respectively. In both cases three beam SIM was used, requiring five phases per direction. The final image reconstruction was performed by applying the locally available Matlab-based code (20).

## Structured illumination microscopy briefly revisited

The image spectrum 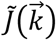 is given by the object 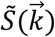 convolved with the illumination spectra multiplied with the transfer characteristics of the imaging system (“optical transfer function”, OTF or system OTF). For the uniform illumination case this is illustrated in fig. 1a. Yet, in SIM the illumination spectrum consists out of distinct delta peaks, typically created by interference of multiple laser beams. In case of two beam interference (two-beam SIM, sinusoidal lateral pattern, no axial modulation) three peaks are located at 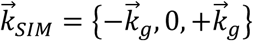. The grating vector 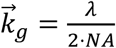 depends on the relative interference angle of the two beams. For three interfering beams, (three-beam SIM, sinusoidal lateral and axial modulation) five peaks are found at 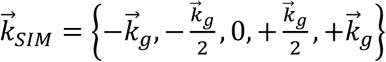.

**Fig. 1:**
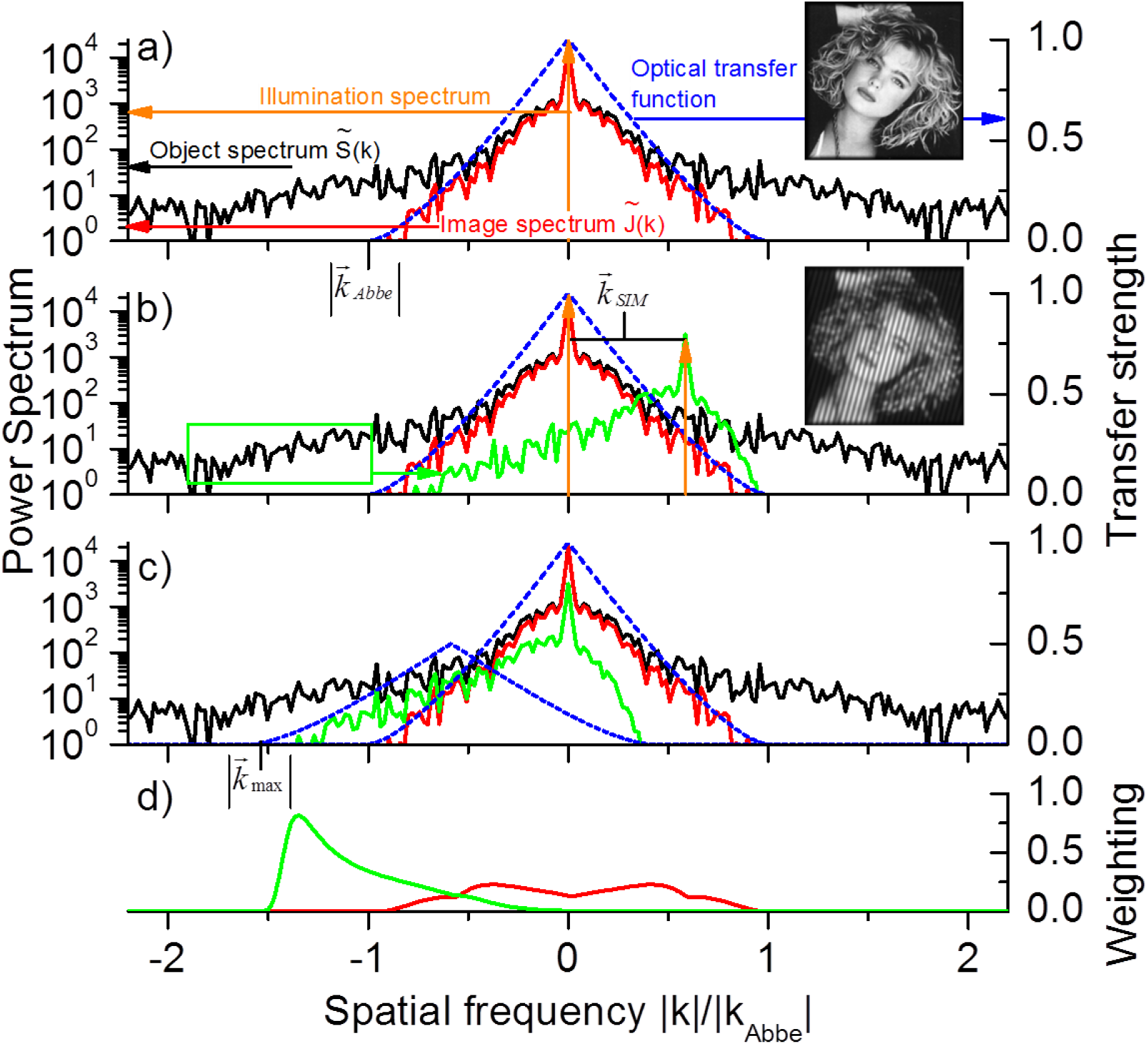
Illustration of the SIM method: a) The frequency transfer of the object spectrum (black) under uniform illumination (orange: illumination spectrum) is governed by Optical Transfer Function (OTF, in blue) and its passband is limited to k_Abbe_ (red), b) Using structured illumination, higher frequency components are modulated into the passband of the OTF (green, note: for sake of clear arrangement only one positive order is shown here), c) After separation and shifting of the orders, the frequency transfer is extended to k_max_, d) The weights which are used for recombining the orders (green: 1^st^ order, red: 0^th^ order)

The recorded image spectrum is the sum of the object spectrum shifted along all elements in 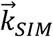 multiplied with the OTF (fig. 1b, for two beam SIM). Consequently, higher object spatial frequencies up to 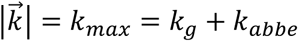 are shifted into the passband of the OTF, leading to the resolution enhancement. The mathematical background of this effect has well been investigated (2-4, 17, 20).

The high-frequency object information (i.e. fine features) is typically retrieved by recording a set of images illuminated using a light pattern with varying phase and applying an unmixing algorithm (22), leading to separated SIM orders. The number of required images is given by the number of elements in 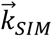 (17). Additionally this is repeated for several (typically three) orientations of the grating in order to achieve a near-isotropic resolution enhancement along the lateral directions x and y. The extended (super-resolved) image spectrum is reconstructed by shifting the unmixed object-components (i.e by 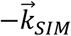) such that their zero object frequencies line up at the center and combining them (Fig. 1c). This is done by a frequency-dependent weighted sum of the individual aligned components. The coefficients are chosen in a way to keep the noise level constant in the reconstructed SIM image (20, 22, 23). The weighting coefficients for -1^st^ and 0^th^ SIM order are sketched in fig. 1d.

The reconstructed spectrum 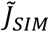 can then be written as a product of the object spectrum and an effective optical transfer function (SIM OTF, c.f. Fig. 1d) 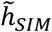 (c.f. supplementary material, sect 2.):

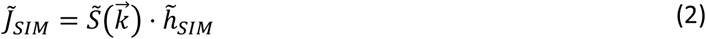

## Important parameters to consider for SIM reconstruction

The accurate determination of **grating parameters** (i.e. the grating phases and their exact position in the Fourier space) is crucial to ensure a proper unmixing and shifting of the SIM orders. They can be evaluated from the raw images with sub-pixel accuracy by exploiting auto- or cross-correlation techniques (17, 22, 24-26). In most available reconstruction software packages this step is automated (27, 28). Furthermore, artifacts related to wrong grating parameter estimation have been classified comprehensively (20, 29).

The **reconstruction parameters**, however, can so far not be retrieved from the raw images. They are classified into parameters related to experimental imperfections and the regularization of the image deconvolution (i.e. solving (2) for 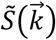). Particularly the latter strongly rely on the signal-to-noise ratio requiring careful readjustment for every image set. The reconstruction parameters are:

### Wiener parameter

To optimize the appearance of the image, a final deconvolution step is necessary, i.e. eq. (1) has to be solved for 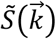. In the presence of noise, a simple division by 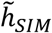 will cause an unacceptable amplification of the noise in frequencies for which 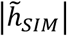 is small. This applies close to the resolution limit. By applying Wiener-filtering a good trade-off between resolution enhancement and noise amplification can be found (c.f. supplementary information, sect 1.). Assuming both, a constant noise as well as a constant object power spectrum, the Wiener-filter is characterized by a constant value *w* (the Wiener parameter). *w* is defined by the inverse signal-to-noise ratio of the image (30). As the latter is, however, generally unknown, the value has to be chosen empirically (22). In the reconstruction process *w* is directly incorporated into the weighting parameters.

### Apodization

The assumption of a constant object spectrum is generally violated. Hence, efficient noise suppression by the Wiener-filter causes a significant distortion of the deconvolved spectrum. An additional radial frequency weighting of the deconvolved spectrum allows further suppressing of high frequency spectral components leaving low frequency components unchanged. Different shapes for the apodization function were suggested such as and exponentially weighted distance transform (22) or a Luckosz bound, which should additionally ensure a non-negative point spread function (31, 32).

### Central frequency suppression

At the detection zero frequency, each SIM order is intentionally suppressed during the reconstruction process due to the high transfer strength of the OTF at this point, being prone to artifacts. In addition, this suppression also enhances the optical sectioning performance, as the detection zero frequency lacks axial frequency support (missing-cone problem), compared to other shifted object components. The latter are more strongly weighted in the presence of efficient zero order suppression. Details on this procedure are found in the supplementary material, sect. 3. The suppression is typically performed by multiplying the order weight with one minus a Gaussian of maximal amplitude A (0 to 1) and variance σ (22).

### Modification parameter of the systems optical transfer function

A disagreement between the simulated system OTF used for reconstruction and the one actually present is compensated by applying an ad-hoc way of modifying the former. To this aim, the simulated (real-valued) transfer function is for instance simply raised to an exponent *γ* slightly different than one (c.f. supplementary material, sect. 4). In effect, this serves to emulate the reduced transfer strength at higher spatial frequencies as, for example, caused by spherical aberrations (33).

An additional parameter allows to account for a decrease in the experimentally achieved modulation depth compared to the theoretical one (c.f. supplementary material, sect. 2). This is sometimes achieved by modifying the strength before summing the various orders.

## RESULTS

### Artifacts in SIM reconstruction, their relation to the image spectrum and to reconstruction parameters

A wrong choice of reconstruction parameters is often easily spotted in the spectrum of the reconstructed data. To illustrate the correlation between the reconstructed data and its spectrum in the presence of typical artifacts arising from poor reconstruction parameter adjustment, the commercial Argolight slide has been used as test sample (Figure 2). Its image and spectrum in the uniformly illuminated case and after deconvolution using the system OTF, respectively, are depicted in figs. 2 a – d. In the spectra the maximum transferred frequency is indicated by the blue circle, which is not changed by the image deconvolution. The latter higher spatial frequencies are, however, more strongly pronounced, so that the image appears sharper.

**Fig. 2:**
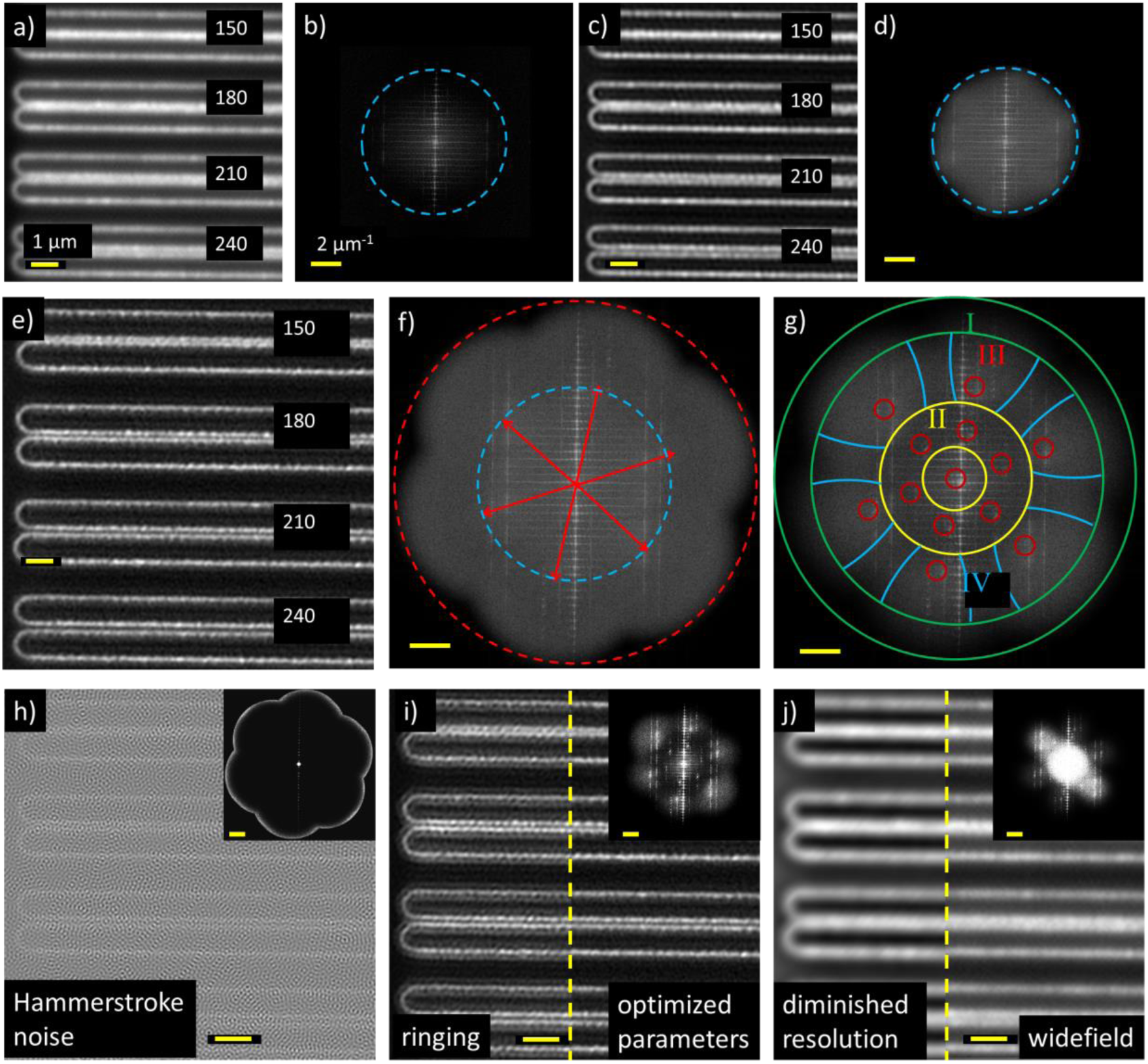
Correlation between a reconstructed image and its spectral magnitude using a test-sample (Argolight slide): a, b) Uniformly illuminated image and spectrum, the blue circle indicates the maximum frequency transfer (k_Abbe_); c, d) like (a) after deconvolution; e, f) SIM reconstruction and its spectral magnitude using well-chosen parameters (the red lines indicate the shifting vectors; red circle indicates maximum frequency transfer k_max_ after reconstruction, blue circle like in (b) and (d)) g) typical deficiency areas in the spectrum (I – IV, c.f. text); h, i, j) representative reconstruction artifacts (e: Hammerstroke noise, f: Ringing, g: diminished resolution) and their spectra (inlays, yellow circle: extended frequency transfer k _Max_), The scale bars for all real space images are 1 µm and for all spectra 2 µm^-1^

In contrast, figs. 2e and f show the image and its spectrum after successful SIM reconstruction. Higher frequencies are transferred, indicated by the red circle, causing the resolution enhancement. The possible resulting artifacts of poor parameter choices are overemphasized in figs. 2 h-I, and it can be summarized in three categories: “Hammerstroke” noise (fig. 2h); ringing (fig. 2i); and, decline in resolution (fig. 2j)

The spectra of images suffering from different artifacts are strongly deviating (inlets in figs 2 h – i). Thus, it is worthwhile to have a closer look at the reconstructed image spectrum in the presence of slightly detuned parameters (fig. 2g), allowing to identify four areas which can be selected for correlated artifacts:

I. <u>Poor transfer of high frequencies (Green)</u> A too strong amplification of the image spectrum at high spatial frequencies will strongly increase high-frequency noise components, since object information in this spectral range is typically low. For frequencies above the resolution limit, on the other hand nothing is transferred, since 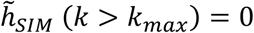. In the real space image, the narrow-band noise amplification is observed by the Hammerstroke pattern. A too strong suppression of high frequency spectral components on the other hand leads to a decrease in resolution.
II. <u>Poor transfer of intermediate frequencies (yellow)</u> If the transfer of spectral components in the intermediate frequency band is suppressed, ringing will be observed in the reconstructed image (c.f. supplementary material, sect. 5, fig S5). The ringing does not show any predominant direction.
III. <u>Residual zero order peaks (red)</u> The weighted recombination might cause an overemphasized transfer of each orders zero detection-frequency. On the one hand this leads to a reduced optical sectioning due to poor axial frequency support for the central order. For higher orders, on the other hand (e.g at 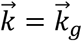) the consequence is residual ringing at the shifting frequency (e.g at 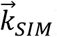) with a predominant direction. This is particularly strong, if filaments are observed which are oriented perpendicularly with respect to the direction of 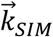 (c.f. supplementary material, sect. 5, fig. S6). Zero-order peaks are particularly strong in the presence of large background signal or relatively thick stained fluorescent samples.
IV. <u>Azimuthally inhomogeneous frequency transfer (blue):</u> An azimuthally inhomogeneous frequency transfer results in a direction-dependent resolution enhancement or weakening, respectively (c.f. supplementary material, sect. 5, fig. S7).

The effects of artifacts in the SIM reconstruction and its spectrum for a biological sample are seen in fig. 3. The resolution enhancement is clearly visible when comparing the SIM image with the uniformly illuminated or deconvolved image (using the system OTF, fig. 3a). For poorly chosen parameters, such as a too large Wiener filter parameter or a too strong apodization no resolution enhancement might be achieved (fig. 3b), due to high frequency transfer suppression. A too small Wiener filter parameter causes an insufficient suppression of noise at frequencies at which the OTF transfer is low. As these frequencies are typically found in a small band around high frequencies close to the resolution threshold (i.e. area I in fig. 2g, c.f. supplementary material fig. S1 and sect. 1), that results in Hammerstroke noise.

**Fig 3:**
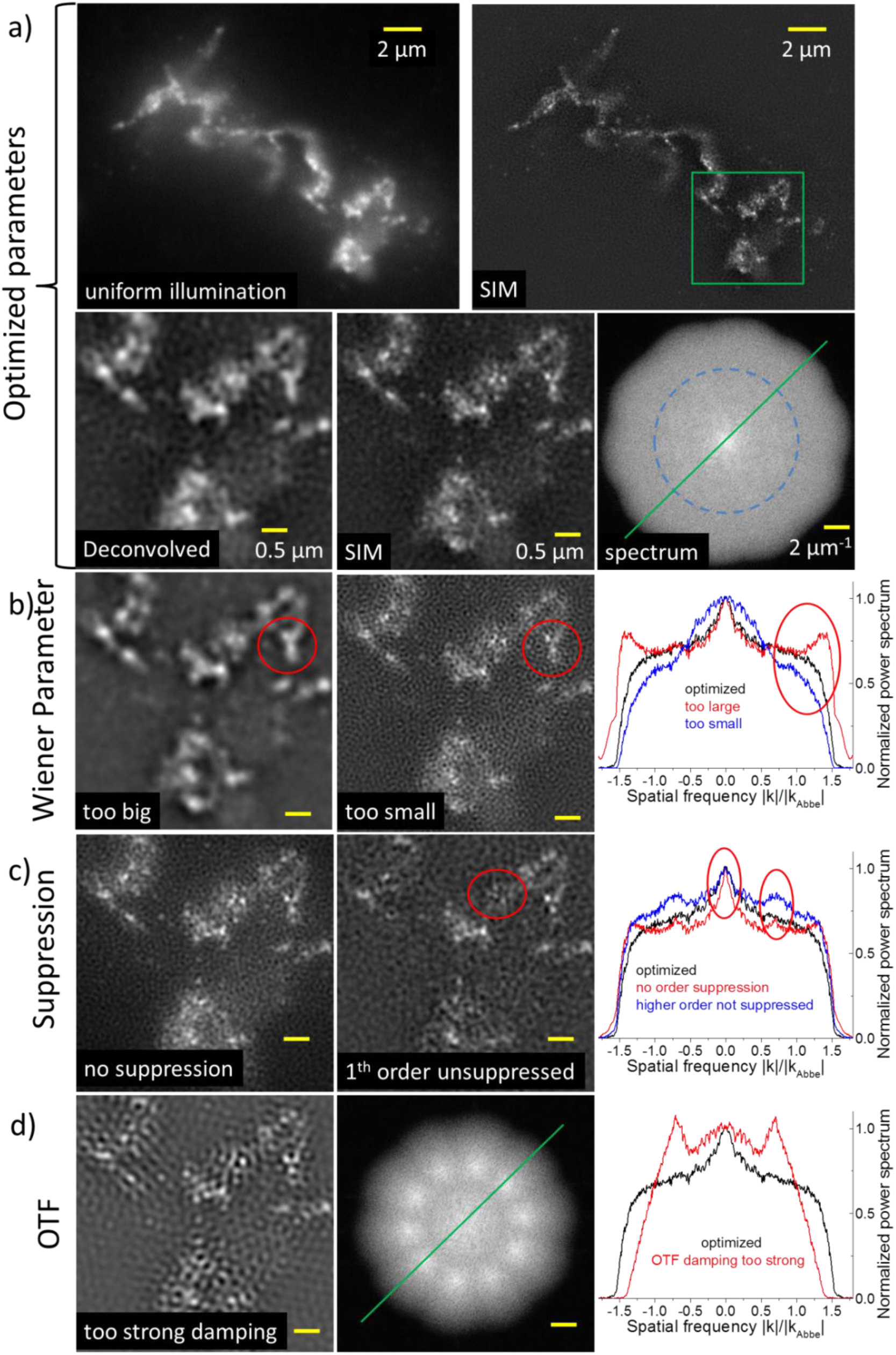
Illustration of the correlation between image artifacts and the reconstruction parameters using a GalNAc-T1 sample: a) Image sets (uniformly illuminated, reconstructed, deconvolved and spectral magnitude) using optimized parameters, the extract correspond to the region marked by the green box in the SIM image, the green line in the spectrum represents the direction of the linecuts (below) and the blue circle k_Abbe_, b) differently chosen Wiener parameter (image extract and linecut through spectrum), red circles: artifact afflicted region, c) different zero order suppression, c) poor OTF damping

The absence of zero-order central frequency suppression leads to an overemphasized transfer of distinct frequencies at 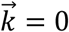 or 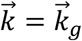 (c.f. fig. 3c) and results in ringing, amplification of residual background and poor sectioning. It is noted, that destructive interference of vicinal ringing structures can cause a loss in information (c.f. red mark in fig. 3c). In case of using many SIM-directions the peaks can fuse into a ring, shaping a frequency band with amplified transfer. In this case the ringing does not show a predominant direction.

A wrong compensation of the theoretical OTF can be an additional reason for ringing. E.g. if the compensating exponent *γ* is chosen to large the wings of the OTF are depleted too strongly. This results in a decrease of the intermediate frequency transfer (fig. 3d), azimuthal inhomogeneities, and a declined resolution as it reduces the maximum transferred object frequency.

In disordered structures (e.g. biological samples), interference of the ringing resulting from vicinal features appears as a statistical pattern and thus is prone to be confused with smaller frequency Hammerstroke noise (i.e. larger dints, c.f. fig. 3b and c). Consequently, reliable artifact identification is tough in the real space image and should be performed by observing the spectrum.

The correlation between different reconstruction parameters and artifacts is summarized in table 1. Note that the parameters are generally not independent, due to the weighted recombination of separated SIM orders. Hence, a change in one parameter may require the adjustment of others to achieve an optimal result. E.g. a strong and broad suppression of the highest SIM-order can cause a homogeneous depletion of high frequencies, thus causing a reduction in resolution. Table 1 depicts the main correlations between artifacts and parameters.

**Tab. 1:** Correlation between artifact and reconstruction parameters

|  | Hammerstroke | Ringing | Resolution diminishing |
| --- | --- | --- | --- |
| Wiener | too small | -- | too large |
| Apodization |  |  |  |
| OTF damping | -- | wrong choice | wrong choice |
| OTF order strength |  |  |  |
| Zero order suppression | too weak (background) | -- | -- |
| higher order suppression | -- | too weak | -- |

### A roadmap for successful reconstruction parameter tuning

Successful image reconstruction requires achieving a homogeneous spectrum of the reconstructed image. The parameter adjustment should be performed by critical investigation of the spectral magnitude. Particularly the tuning range of the Optical Transfer Function (OTF) related parameters can be vastly reduced by proper characterization of the experiment and preprocessing of the data:

- Measure the PSF of the system and compute the OTF by a Fourier transform.
- Measure the power of the individual laser beams contributing to the grating and calculate the amplitudes
- Use appropriate cover glass thicknesses (typically high precision cover glass thicknesses of 170 ± 5 µm) and suited immersion oil to minimize spherical aberrations
- Do a proper background subtraction of the raw data.

For the optimization of the processing it is recommended to start with the following parameter settings:

- Use a very low Wiener parameter (*w*^2^ ≈ 10^−10^). This will cause a tremendous over rating of high frequencies, but it will directly show the maximum achievable frequency.
- No apodization
- Strong zero-frequency suppression (complete suppression for all orders)
- Use either the experimental OTF (if possible) or a slightly modified theoretical OTF (in case of an exponential damping, use an exponent around 1.1)
- If possible, set the amplitudes of the interfering laser beams as measured during the experimental preparation (c.f. above) in the reconstruction package.

Continue with the following iterative optimization process:

- Reduce the Wiener parameter until high frequency components are not over rated anymore. Typical values rang from (10^−5^ < *w*^2^ < 10^−8^).
- Azimuthally diminished spectrum and too weakly transferred small frequencies: reduce the OTF damping and adjust the OTF – order strength parameter (if possible)
- Distinct peaks at zero as well as shifting frequencies: Adjust respective order suppression strength and width. The suppression should be in the range between 98 % and 100 %
- Optimize the Wiener parameter and the apodization in order to extend the spectrum as far as possible without over rating high frequencies. The apodization should be fairly small.
- Repeat the steps iteratively until a homogeneous spectrum is reached.

### Implementation of the parameters in selected SIM reconstruction packages

The reconstruction parameters are differently implemented in the available software packages as summarized in tab 2. The open SIM toolbox, which is a Matlab package as well, is not considered in this work, as it only allows to reconstruct two –beam SIM images.

**Tab. 2:**
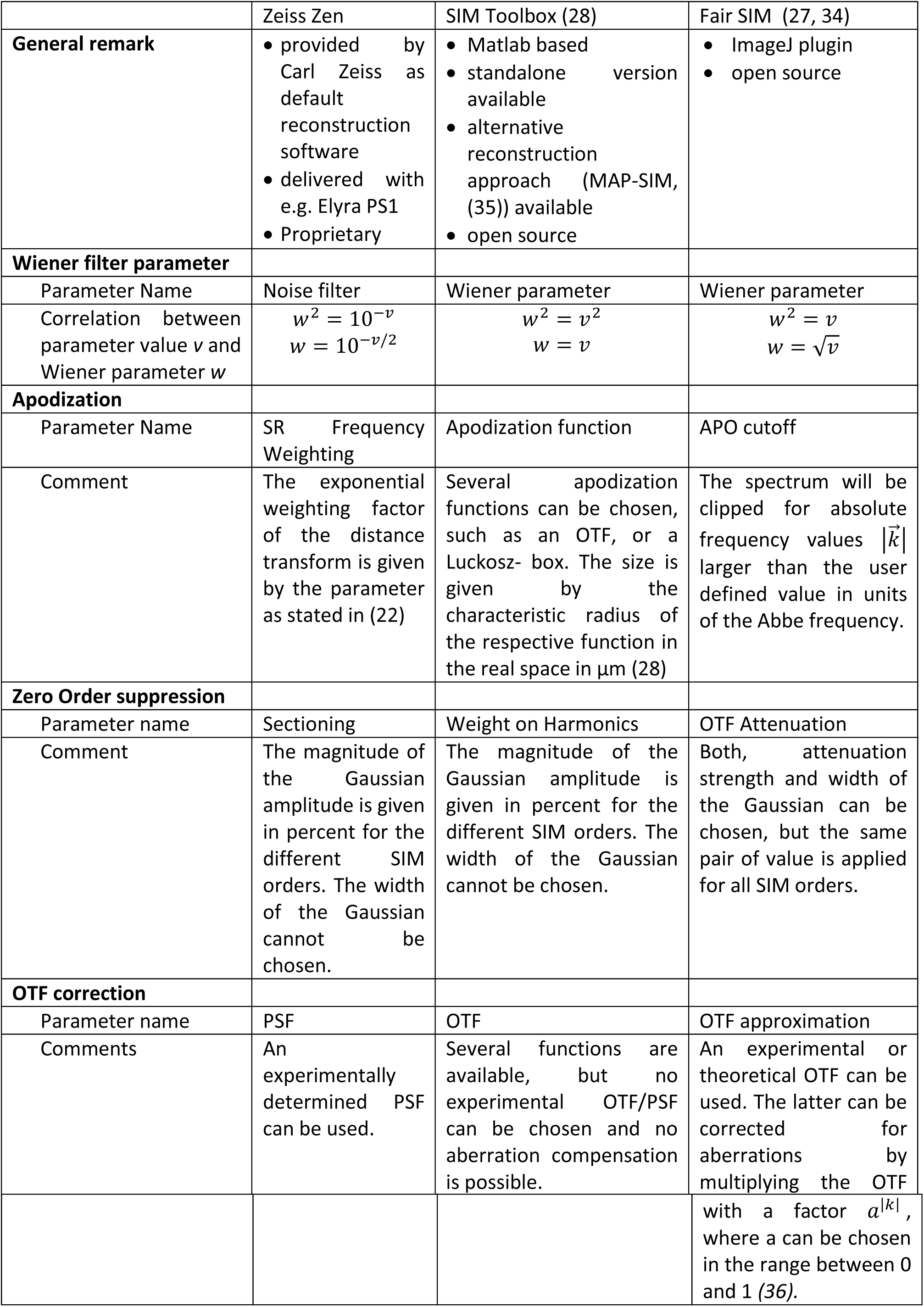
Comparison of the parameter implementation in different available software packages

## Conclusion

In this work the impact of reconstruction parameters in SIM was analyzed. The parameters comprise the Wiener filter parameter, the apodization function, zero-frequency suppression and modifications of the optical transfer function. They are particularly critical, as they have to be optimized for each individual reconstruction.

The artifacts resulting from poor parameter selection can be classified into ringing, amplification of Hammerstroke noise and reduction of resolution. Whereas reduced resolution and Hammerstroke noise are mostly caused by a poorly chosen wiener parameter and/or apodization, a wrong correction of the OTF or insufficient zero-order suppression lead to ringing. Distinguishing between the different artifact types in the real image is, however, difficult and can lead to misinterpretation. Instead the reconstructed spectrum should be observed during the optimization process. A homogeneous spectrum is suggested to be the goal of the optimization. A roadmap for efficient and successful parameter adjustment is provided. Additionally, different available software packages for SIM reconstruction were compared.

## Acknowledgement

Funding by the German research foundation (Deutsche Forschungsgesellschaft, DFG) in the framework of the Project SFB TR166 “Receptorlight”, TP B05 as well as by the “Thüringer Aufbaubank” (TAB) in the framework of the Inphodeg project is gratefully acknowledged. The Swedish Foundation for Strategic Research (SSF) for founding the Research Infrastructure Fellow program no. RIF14-0079. The COST Action (CA15124) NEUBIAS for supporting Maria Smedh with a Short-Term Scientific Mission. Furthermore, the authors thanks Arnaud Royon and the Argolight Company (Pessac, Bordeaux, France) for providing the ArgoSIM slide.

